## Supplementary material for "Optimizing information transmission in neural induction constrains cell surface contacts of ascidian embryos"

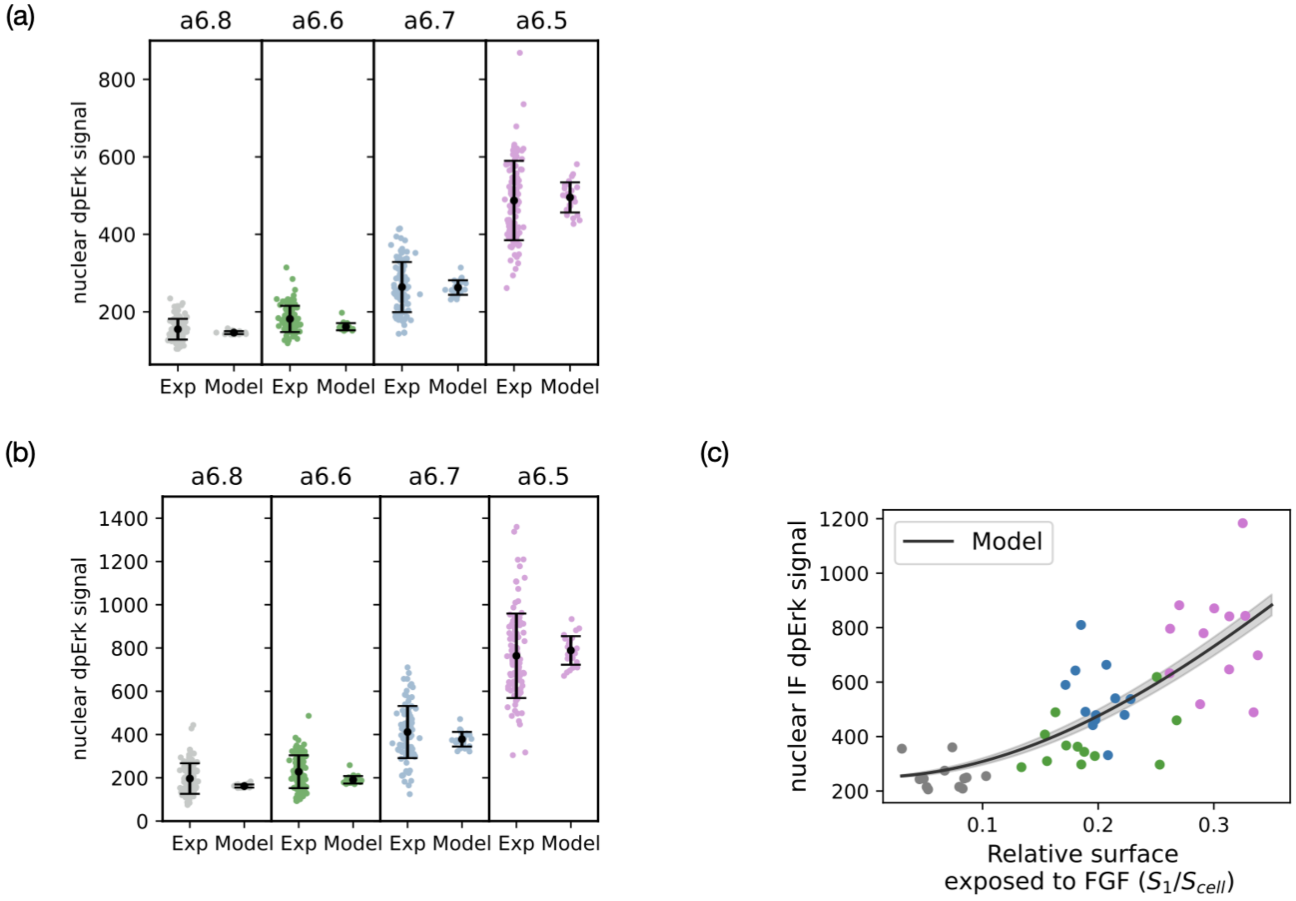


FIG. S1. (a) Nuclear dpERK immunofluorescence (IF) signals in the a6.5, a6.6, a6.7 and a6.8 cell types as measured in IF experiments in wild type embryos (left) and computed with the model (right). Each point represents a single cell and modeling results are computed using the measured values of S1. Means and standard deviations are shown in black. The parameter values used to reproduce the experimental data are listed in table I. $A = 5000$, $B = 140$. (b) Nuclear dpERK immunofluorescence (IF) signals in the a6.5, a6.6, a6.7 and a6.8 cell types as measured in IF experiments in embryos in which the ephrin pathway was inhibited (left) and computed with the model (right). Each point represents a single cell and modeling results are computed using the measured values of S1. Means and standard deviations are shown in black. The parameter values used to reproduce the experimental data are listed in table I. $A = 8000$, $B = 150$. To mimic the absence of ephrin, we imposed $e = 10^{-5}$. (c) Experimental data in the absence of ephrin. The plot shows the nuclear dpERK immunofluorescence (IF) signals measured experimentally in individual a-line cells as a function of the relative area of cell surface contact with FGF-expressing cells ($S_1/S_{cell}$). Experimental data obtained in the different cell types are shown with dots of different colors: a6.8 cells are shown in gray, a6.6 cells in green, a6.7 cells in blue, and a6.5 cells in magenta. The experimental data obtained in embryos in which the ephrin pathway is inhibited are compared with the model predictions (shown with a black line). The shaded region represents the noise in the level of ERK* fluorescence predicted by the model using the Langevin approach. The model predictions are obtained as described in section III A, V A and V B. The parameter values used to reproduce the experimental data are listed in table I. $A = 10000$, $B = 250$. To mimic the absence of ephrin, we imposed $e = 10^{-5}$.

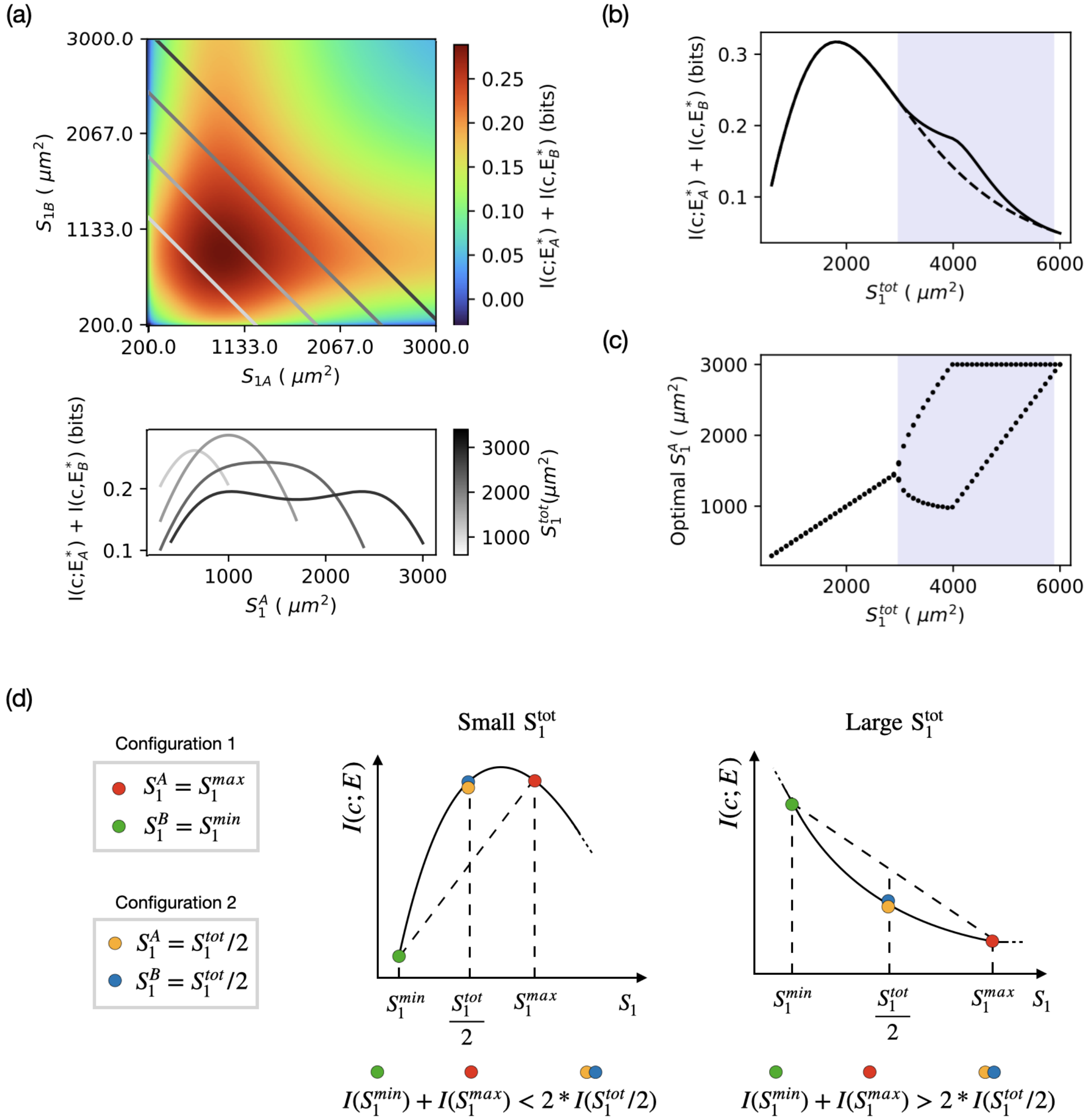


FIG. S2. (a) Independent cell case. The heat map represents the information transmitted assuming the two cells are independent $I = I(c; E_A) + I(c; E_B)$ as a function of the area of cell surfaces exposed to FGF ($S_1^A$ and $S_1^B$). The presence of the constraint restricts the possible values of $S_1^A$ and $S_1^B$ to straight lines defined by $S_1^B = S_1^{\text{tot}} - S_1^A$. The straight lines colored with different shades of gray represent the accessible values of $S_1^A$ and $S_1^B$ for the values of $S_1^{tot}$ used in the plot below. The plot below represents $I = I(c; E_A) + I(c; E_B)$ as a function of $S_1^A$ for different values of the constraint $S_1^{\text{tot}}$ (see colorbar). (b) Maximal information (solid line, obtained when the two cells expose the optimal surface areas to FGF) and the information transmitted when the two cells have the same surface areas exposed to FGF (dashed line) as a function of $S_1^{\text{tot}}$. (c) Optimal values of $S_1^A$ as a function of $S_1^{\text{tot}}$. The purple background highlights the regions of the plots where it is maximally informative for the two cells to expose different surface areas to FGF. Panels (a), (b) and (c) are obtained assuming $\mu_c = 40$, $\sigma_c = 1$, $S_{\text{cell}}^A = S_{\text{cell}}^B = 6000\ \mu m^2$. (d) Scheme illustrating why the optimal geometrical configuration of the cells vary with $S_1^{\text{tot}}$. We compare two geometrical configurations of the cells: configuration 1 in which $S_1^A = S_1^{max}$ and $S_1^B = S_1^{min}$, and configuration 2 in which $S_1^A = S_1^B = S_1^{\text{tot}}/2$. $S_1^{max}$ and $S_1^{min}$ are the maximal and minimal possible values for $S_1$ (fixed by the values of $S_{cell}$ and $S_1^{\text{tot}}$). At low values of $S_1^{tot}$ the part of the curve accessible is concave. In this condition, it is easy to see that the configuration of the cells that maximizes information transmission is configuration 2 ($S_1^A = S_1^B = S_1^{tot}/2$). At large values of $S_1^{tot}$, the part of the curve that is accessible is convex. In this condition, the configuration that maximizes information transmission is configuration 1 ($S_1^A \neq S_1^B$).

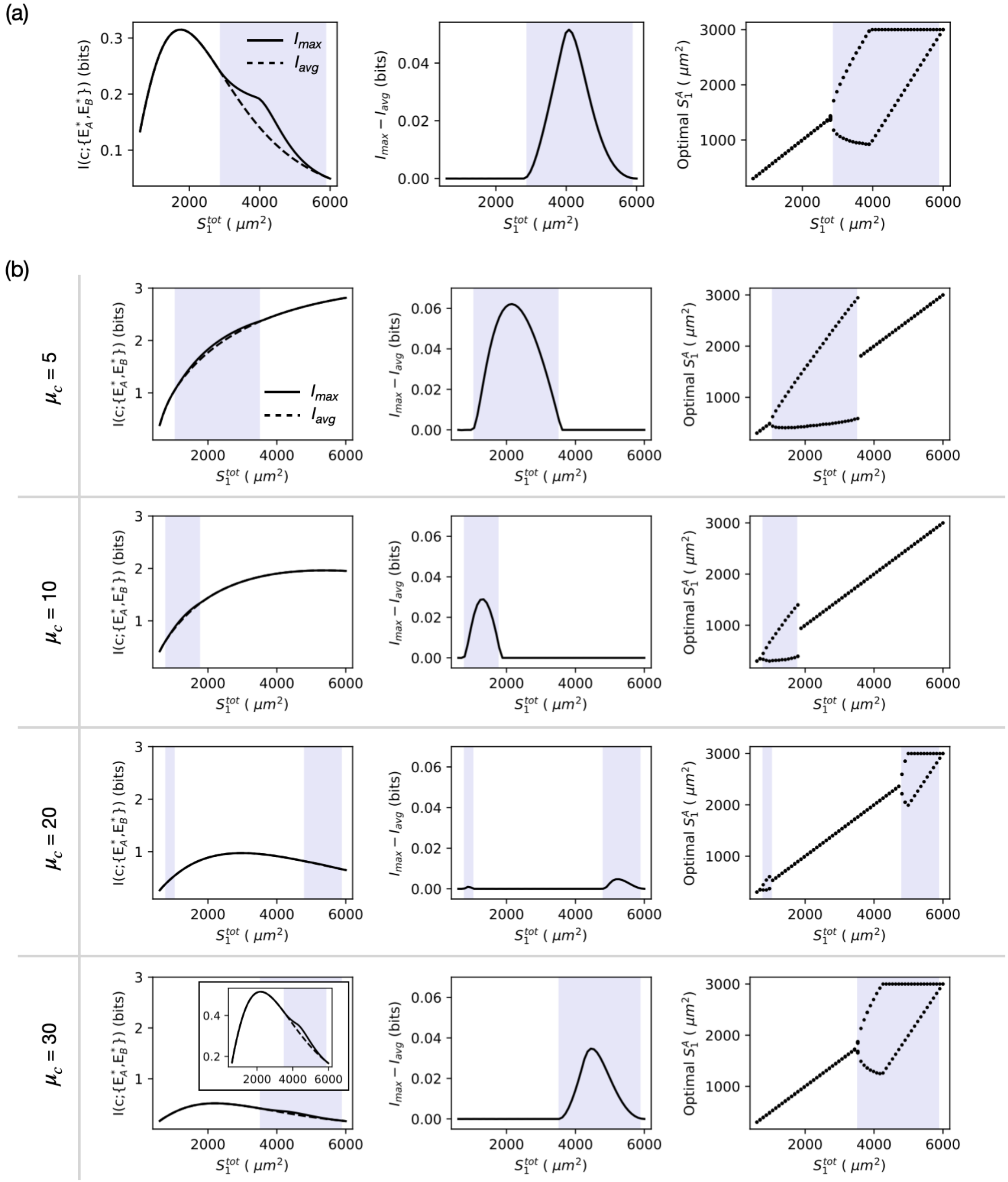


FIG. S3. (a) Information transmission in the absence of ephrin. The plot on the left shows the maximal information ($I_{\max}$) transmitted between $c$ and $\{E_A^*, E_B^*\}$ (solid line, obtained when the cells expose the optimal surfaces to FGF) and the information transmitted when the two cells expose the same surface to FGF ($I_{\text{avg}}$, dashed line) as a function of $S_1^{\text{tot}}$. The plot in the middle shows the difference between $I_{\max}$ and $I_{\text{avg}}$ as a function of $S_1^{\text{tot}}$. The plot on the right shows the values of the optimal surface $S_1^{A*}$ as a function of $S_1^{tot}$. The purple background highlights the regions of the plots where it is better, in order to maximize information transmission, for the two cells to break the symmetry and expose different surfaces to FGF. These plots are obtained assuming that the input distribution $P(c)$ is centered around $\mu_c = 40$ and that the two cells have the same total surface $S_{\text{cell}}^A = S_{\text{cell}}^B = 6000\ \mu m^2$. To model the absence of ephrin we used $e = 10^{-5}$. (b) Influence of the position $\mu_c$ of the input distribution on information transmission. The plots in the first column show the maximal information ($I_{\max}$) transmitted between $c$ and $\{E_A^*, E_B^*\}$ (solid line, obtained when the cells expose the optimal surfaces to FGF) and the information transmitted when the two cells expose the same surface to FGF ($I_{\text{avg}}$, dashed line) as a function of $S_1^{\text{tot}}$. The plots in the second column show the difference between $I_{\max}$ and $I_{\text{avg}}$ as a function of $S_1^{\text{tot}}$. The plots in the third column show the values of the optimal surface $S_1^{A*}$ as a function of $S_1^{\text{tot}}$. The purple background highlights the regions of the plots where it is better, in order to maximize information transmission, for the two cells to break the symmetry and expose different surfaces to FGF. These plots are obtained assuming that the two cells have the same total surface $S_{\text{cell}}^A = S_{\text{cell}}^B = 6000\ \mu m^2$.

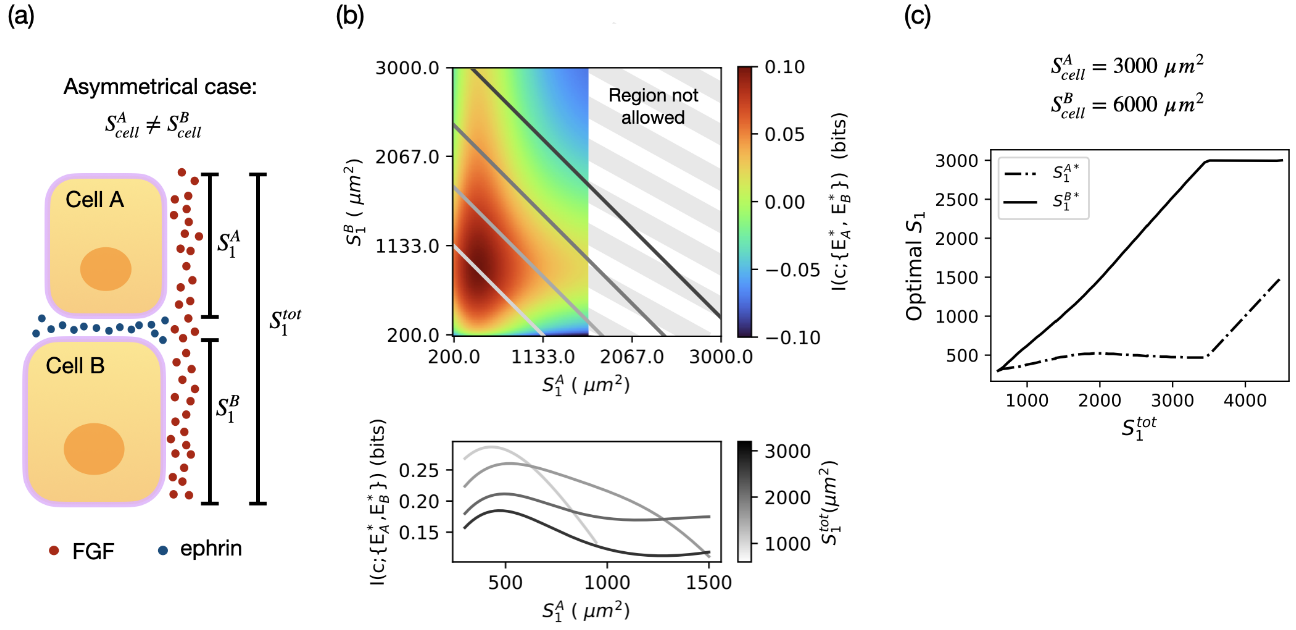


FIG. S4. Two-cells: asymmetrical case. (a) In the asymmetrical case we consider that the two cells have different cell surface areas ($S^A_{cell} \neq S^B_{cell}$). $S^A_1$ and $S^B_1$ are the surface areas of the cells exposed to FGF. $S^A_2$ and $S^B_2$ (not shown in the cartoon) are the surface areas of the cells exposed to ephrin. In each cell, $S^{A,B}_2$ is computed from the corresponding value of $S^{A,B}_1$ using equation Eq.(5). The value of $S_1$ in each cell is constrained to be at most equal to $S_{\rm cell}/2$, with $S_{\rm cell}$ being the total surface area of the cell. $S^{\rm tot}_1$ represents the total surface area where FGF is present. The presence of the constraint implies that $S^A_1 + S^B_1 = S^{\rm tot}_1$. (b) The heatmap represents the information transmitted between the FGF input $c$ and the ERK output $\{E^*_A, E^*_B\}$ as a function of the area of cell surfaces exposed to FGF ($S^A_1$ and $S^B_1$). The presence of the constraint restricts the possible values of $S^A_1$ and $S^B_1$ to straight lines defined by $S^B_1 = S^{\rm tot}_1 - S^A_1$. The straight lines colored with different shades of gray represent the accessible values of $S^A_1$ and $S^B_1$ for the values of $S^{tot}_1$ used in the plot below. The plot below represents $I(c; \{E^*_A, E^*_B\})$ as a function of $S^A_1$ for different values of the constraint $S^{\rm tot}_1$ (see colorbar). (c) The plot shows the optimal values of the surfaces exposed to FGF ($S^{A*}_1$ with a solid line and $S^{B*}_1$ with a dashed line) as a function of the value of the constraint $S^{tot}_1$. The optimal values are obtained maximizing information transmission assuming $\mu_c = 40$, $S^A_{cell} = 3000\ \mu m^2, S^B_{cell} = 6000\ \mu m^2$.
